# Distal inhibition is recruited for action but not sensation in the primate superior colliculus

**DOI:** 10.64898/2026.11.11.765869

**Authors:** Clara Bourrelly, Feiran Yang, Neeraj J. Gandhi

## Abstract

Sensorimotor transformation requires neural circuits to coordinate excitation and inhibition, but how and whether this organization changes as the circuit transitions from processing sensation to action remain unresolved. We recorded laminar local field potentials across superior colliculus (SC) layers in four male macaques and computed current source density to resolve transmembrane currents during saccades to a target at one of eight isoradial directions. During the visual response, excitation was spatially confined to the receptive field, with little modulation at distant sites. During saccade generation, local excitation remained strongest for movements toward the receptive field, while inhibition extended across the SC map, including locations where spiking remained at baseline. Because the same recordings detected distal inhibition during movement, its absence during vision cannot reflect insufficient sensitivity, indicating that distal inhibition is selectively recruited during action. After saccade completion, excitation emerged regardless of saccade direction, potentially reflecting a rebound from, or release of, the preceding inhibition. These patterns were only partly reflected in spiking activity, which primarily captured local excitation, unmasking circuit dynamics largely invisible at the level of neuronal output. Together, our findings reveal a striking reconfiguration of SC population dynamics across sensorimotor transformation, spanning from spatially restricted excitation during sensation, to local excitation coupled with distal inhibition during movement generation, then to global post-movement excitation. Thus, fixed circuit architecture can express a sequence of behaviorally gated excitation-inhibition states largely invisible to spiking output.

## Introduction

The brain transforms fleeting sensory inputs into motor commands. This transformation requires neural circuits to dynamically coordinate excitation and inhibition across space and time. Although this principle is central to systems neuroscience, how excitatory and inhibitory interactions reorganize within a circuit as it transitions from sensation to action remains poorly understood. The superior colliculus (SC) provides a uniquely tractable model for addressing this question. As a layered and topographically organized structure essential for visually guided saccades, the SC contains sensory, visuomotor, and motor neurons arranged along a shared retinotopic map (Basso & May 2017, Gandhi & Katnani 2011, Hafed et al 2023, May 2006, Wurtz & Albano 1980). SC population activity during both stimulus processing and saccade generation forms a localized “mound” centered on the target’s spatial location (Anderson et al 1998), a profile thought to arise from structured interactions between excitation and inhibition within and across SC layers (Arai et al 1994, Munoz & Istvan 1998, Takahashi et al 2010).

Evidence from several approaches supports structured, spatial organization of excitation and inhibition within the SC. Slice studies have revealed center excitation-surround inhibition within the superficial layers, while intermediate layers exhibit more spatially extensive interactions, including both long-range excitation and inhibition (Isa & Hall 2009, Meredith & Ramoa 1998, Phongphanphanee et al 2008, Phongphanphanee et al 2014). *In vivo* studies in cats and monkeys provide additional evidence for interactions between distant SC locations (Hoffmann & Straschill 1971, Mascetti & Arriagada 1981, Munoz & Istvan 1998, Takahashi et al 2010). In behaving primates, visual responses to a stimulus within a neuron’s receptive field are also reduced firing rates when competing stimuli are presented elsewhere, an effect attributed in part to lateral inhibition across the collicular map (Basso & Wurtz 1998, McPeek & Keller 2002). Together, these observations have supported a framework in which local excitation and distal inhibition shape activity across the SC map. Yet this framework has not been tested in behaving animals by directly assessing the spatial organization of the underlying inhibitory activity.

Previous evidence for inhibition has relied largely on reduced preparations, anesthesia, perturbations, or inference from changes in spiking activity (Dorris et al 2007, Haider et al 2013, Jagadisan & Gandhi 2017a, Jagadisan & Gandhi 2016, Jagadisan & Gandhi 2017b, Munoz & Istvan 1998, Populin 2005, Skoblenick & Everling 2012, Taub et al 2013, Vizuete et al 2012). Reliance on spiking is particularly limiting because little or no activity at a given SC location cannot distinguish active inhibition from a lack of excitation. We therefore asked how the spatial organization of excitation and inhibition changes as the SC transitions from visual processing to movement generation and movement completion. We recorded laminar local field potentials in behaving primates and applied current source density (CSD) analysis to estimate net transmembrane currents across SC layers (Freeman & Nicholson 1975, Nicholson & Freeman 1975, Pettersen et al 2006). By systematically varying target location relative to each recording site, we mapped the spatial organization of subthreshold population activity during visual, motor, and post-motor epochs, allowing us to examine whether excitatory and inhibitory interactions maintain a common spatial organization or are reconfigured across behavioral states.

We found that the spatial organization of excitation and inhibition changed markedly across the visuomotor transformation. During visual processing, excitation was confined to the receptive-field location, with no evidence of inhibition at distant sites. During saccade generation, in contrast, the same recording sites showed local excitation accompanied by inhibition extending across the map, including locations where spiking remained at baseline. Crucially, because distal inhibition was detected in these same recordings during movement, its absence during visual processing is unlikely to reflect insufficient sensitivity. Following movement completion, activity reorganized into widespread excitation across target directions, suggestive of circuit resetting. Thus, rather than reflecting a fixed spatial architecture, excitation and inhibition across the SC map are dynamically reconfigured across visual, motor, and post-motor states. These findings highlight the importance of examining subthreshold population dynamics and reveal a state-dependent organization of circuit interactions as the SC transitions from sensation to action.

## Results

To examine neural dynamics across SC layers, we applied current source density (CSD) analysis to laminar local field potential (LFP) recordings obtained while monkeys performed delayed saccade tasks (Fig. 1). The CSD computes the second spatial derivative of extracellular voltage across contacts, providing a spatially resolved estimate of net transmembrane current flow. In this framework, negative deflections (sinks) correspond to net inward current, typically associated with excitatory input, whereas positive deflections (sources) reflect net outward current, often linked to inhibitory input or passive return currents. Although sinks and sources cannot be interpreted as pure excitation and inhibition in a one-to-one manner, CSD analysis allows inference of the laminar distribution and timing of currents that are not directly observable from spiking activity alone. This approach enabled us to track how extracellular voltage evolves across depth and behavioral epoch during the visuomotor transformation.

**Figure 1.**
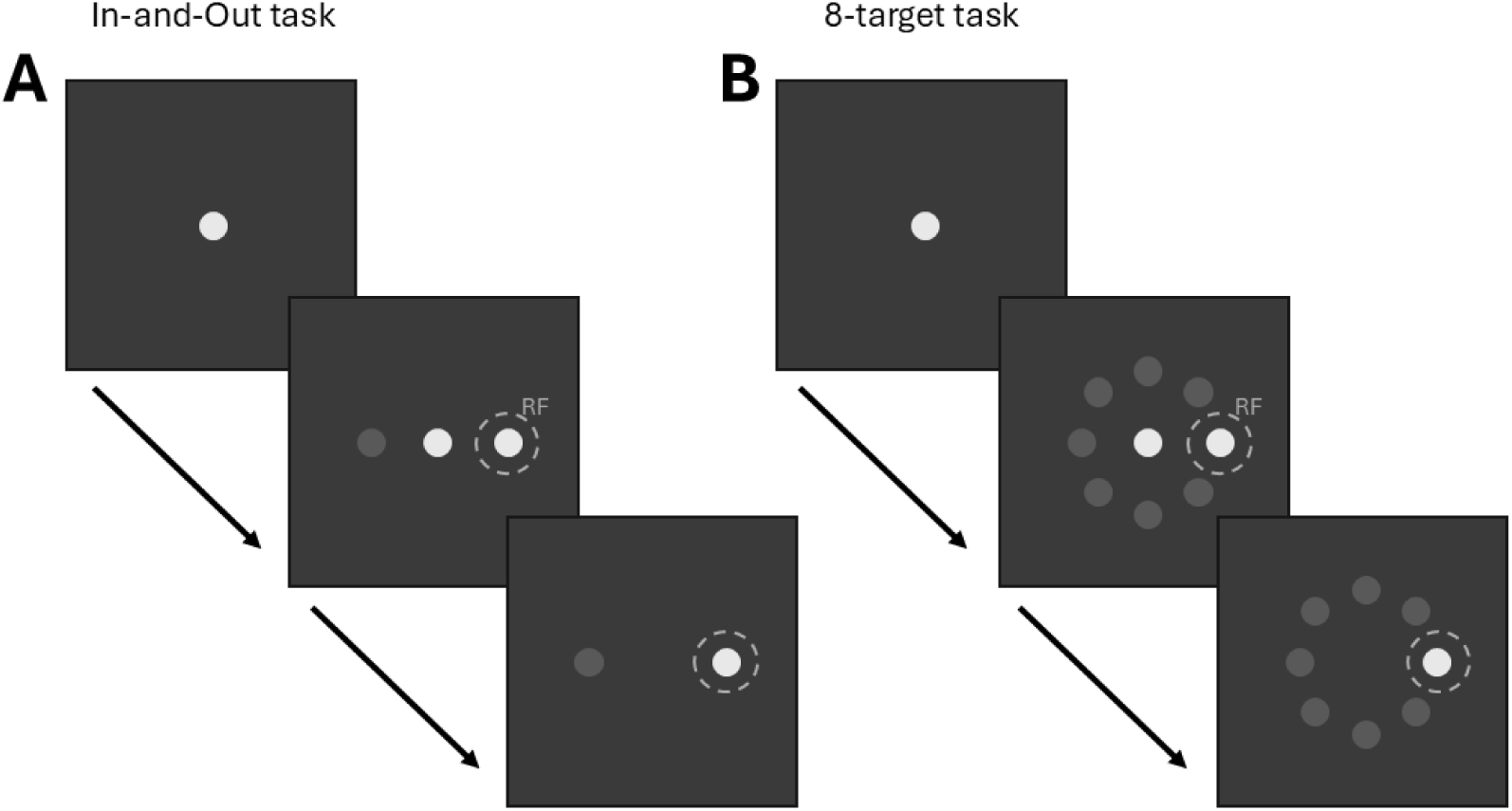
Delayed-saccade task configurations. (A) In-and-Out task. After fixation, a peripheral target appeared either within the receptive field (RF; T1) or at the diametrically opposite location. Following the delay period, the fixation point disappeared, instructing the monkey to make a saccade to the target. (B) Eight-target task. The same delayed-saccade sequence was used, but the target could appear at one of eight locations spaced 45° apart along an imaginary circle. All targets had the same eccentricity from fixation and differed only in direction. For Monkeys SU and DU, the target array was oriented according to the estimated RF location, with T1 positioned within the RF and the remaining targets placed at successive 45° angular increments from T1. For Monkey BL, targets were presented at a fixed eccentricity of 10° at directions ranging from 0° to 315°, and the target closest to the estimated RF location was designated as T1. Dashed circles indicate the RF location.

### Visual activity is spatially localized to the receptive field and superficial layers

Figure 2 shows, for one example session (recorded in Monkey BL), the trial-averaged spike (Fig. 2A,B), LFP (Fig. 2C,D), and CSD (Fig. 2E-H) activity across depth for targets presented inside the receptive field (in-RF) or the diametrically opposite location (out-RF) recorded during the In-and-Out task. Signals are aligned either to target onset or to saccade onset and are displayed across all recording contacts.

**Figure 2.**
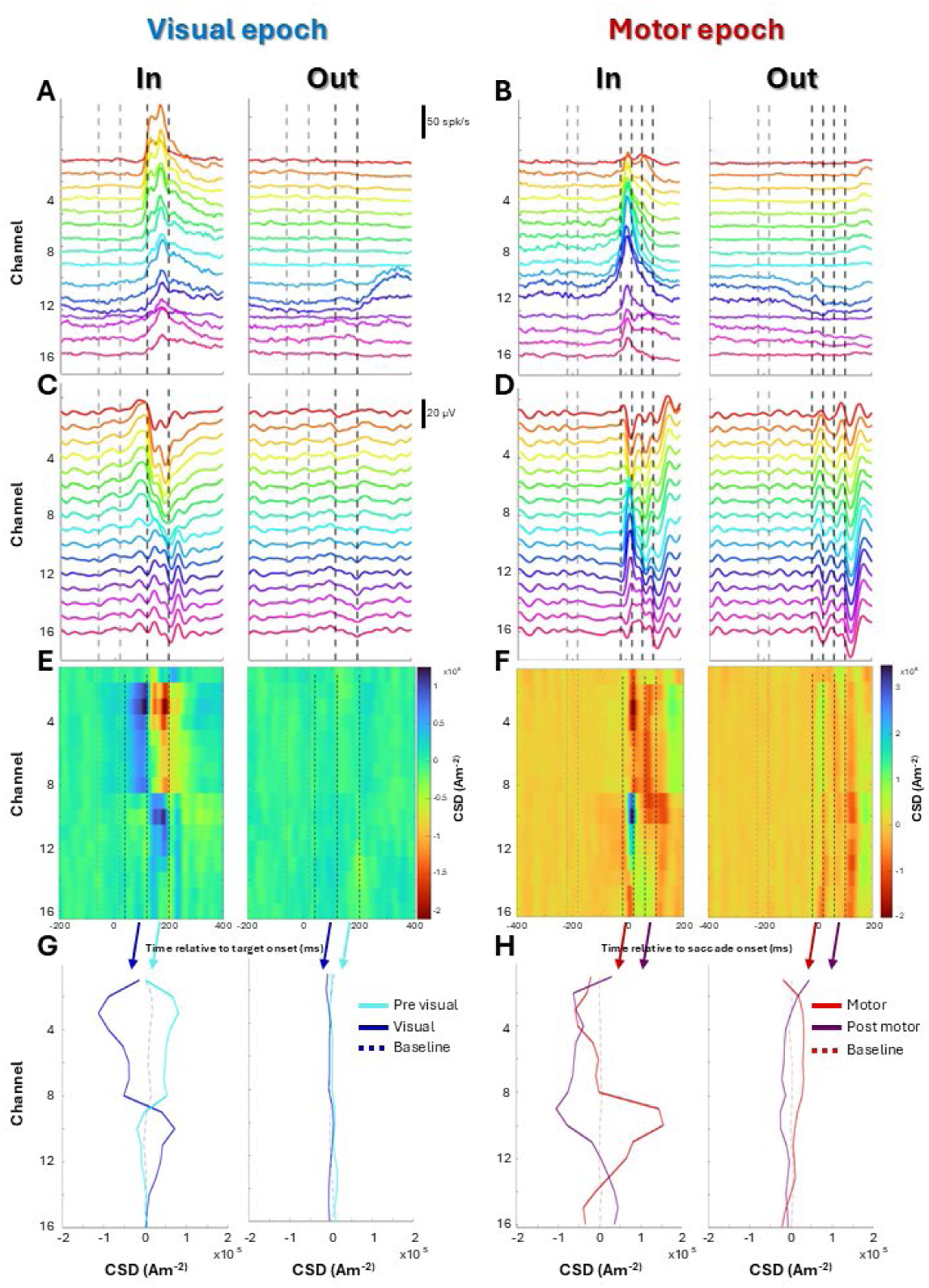
Spike, LFP, and CSD signals during the In-and-Out task from an example dataset. All data are from the same example recording session in Monkey BL. Average spiking activity across depth aligned to target onset (A) and saccade onset (B), shown for trials with targets inside the receptive field (In; left) and outside the receptive field (Out; right). LFP signals aligned to target onset (C) and saccade onset (D), and corresponding CSD signals aligned to target onset (E) and saccade onset (F), are displayed in the same format. (G) Average CSD across depth during the pre-visual (solid cyan), visual (solid blue), and baseline (dashed blue) windows for In-RF (left) and Out-RF (right) trials. (H) Average CSD across depth during the motor (solid red), post-motor (solid purple), and baseline (dashed red) windows for In-RF (left) and Out-RF (right) trials. Traces at the top correspond to superficial channels. Dashed vertical lines indicate analysis time windows.

Following target onset, spike bursts and LFP modulations emerge across depth, reaching their largest amplitude in the superficial layers shortly after the appearance of targets in the RF (Fig. 2 A,C, left). The strongest modulations occur approximately 140-220 ms after target onset (visual response window, black dashed lines). In contrast, activity is markedly weaker for out-RF targets (Fig. 2A,C right), and signals remain close to baseline (gray dashed lines) across most channels.

The CSD signal (Fig. 2E) reveals that targets presented in the RF evoke a prominent current sink (red hotspot in Fig. 2E, left) in superficial layers accompanied by a source (blue hotspot in Fig. 2E, left) in deeper layers, emerging within the same post-stimulus time window. For out-RF targets, these sink-source pattern is absent (Fig. 2E right). Average CSD activity across depth during the baseline (gray) and visual windows (blue) is summarized in Figure 2G, confirming that modulation is robust for in-RF targets and minimal for outRF targets. In this example session, an additional feature is visible in the LFP signal preceding the main LFP trough in superficial channels (Fig. 2C, left), consisting of a transient increase in voltage. This deflection is accompanied by a corresponding current source in the CSD signal (blue hotspot preceding red hotspot in Fig. 2E). This “pre-visual” modulation is evident within a time window of 40-120 ms after target onset, during which the CSD activity mirrors the modulation observed in the visual epoch (Fig. 2G, cyan traces). Notably, this feature was not consistently observed across datasets (present in 28 of 42 sessions), so it was not analyzed systematically and is not considered further, although its potential significance is addressed in the Discussion.

### Motor and post-motor activity extend beyond the receptive field

Trial-averaged spike, LFP, and CSD activity aligned to saccade onset are shown in Figure 2B, D, and F and H, respectively. Three analysis epochs were considered: a motor baseline window preceding movement, a motor window centered on saccade onset, and a post-motor window following movement (see Method for details). Around saccade onset, strong modulations in spiking activity (Fig. 2B, left) and LFP signals (Fig. 2D, left) are observed when saccades are directed toward the RF. These modulations are most prominent across intermediate and deeper channels and peak near saccade onset (i.e., during the motor window). When saccades are directed away from the RF (out-RF condition, right panels), spike bursts are absent (Fig. 2B, right), and LFP modulations are reduced but not negligible (Fig. 2D, right). During the post-motor epoch, spiking activity returns toward baseline for RF saccades (Fig. 2B, left) and remains absent for out-RF saccades (Fig. 2B, right). In contrast, LFP activity remains modulated for a longer period, although the magnitude of modulation is generally larger for in-RF than out-RF trials (Fig. 2D, compare left and right panels). Notably, LFP modulation is observed in out-RF trials even when spiking activity is absent, indicating that subthreshold activity may be present despite the lack of neuronal output.

The corresponding CSD profiles revealed the laminar organization of this neural activity and differed markedly between in-RF and out-RF saccades (Fig. 2F). For saccades directed toward the RF (left), the motor epoch is characterized by a prominent superficial sink (red hotspot) accompanied by a deeper source (blue hotspot) around saccade onset. In contrast, for out-RF saccades (right), the motor epoch exhibits source activity, with no accompanying sink. During the post-motor epoch, a sink is observed for both RF and out-RF trials. Average CSD activity across depth during the motor and post-motor windows is summarized in Figure 2H, confirming that motor-epoch activity differs across conditions, with a superficial sink and deeper source for in-RF saccades and a dominant source for out-RF saccades (red traces), whereas post-motor activity is characterized by a sink in both conditions (purple traces).

### CSD activity differs across visuomotor epochs and RF locations

Figure 3 quantifies CSD activity across depth during the visual (blue), motor (red), and post-motor (purple) epochs for in- and out-RF target locations for all sessions. Panel A shows the average CSD profiles averaged across all monkeys. For in-RF targets, a clear sink is observed during the visual epoch (negative deflections in blue traces), followed by a paired sink and source during the motor epoch (negative deflections in superficial channels and positive deflections in deeper channels in red traces), and a prominent source during the post-motor epoch (positive deflections in purple traces). In contrast, out-RF targets show little or no modulation during the visual epoch (blue traces near zero), a source during the motor epoch (red traces, positive values), and a sink during the post-motor epoch (purple traces, negative values). These patterns are consistent across datasets, both when within and across monkeys, and closely match those observed in the individual example dataset (Figure 2). Note that visual-epoch sinks are more pronounced in individual datasets but appear attenuated in the pooled profiles, likely because alignment on the motor epoch disperses activity across depth and reduces the median amplitude (see individual dataset traces, dashed lines).

**Figure 3.**
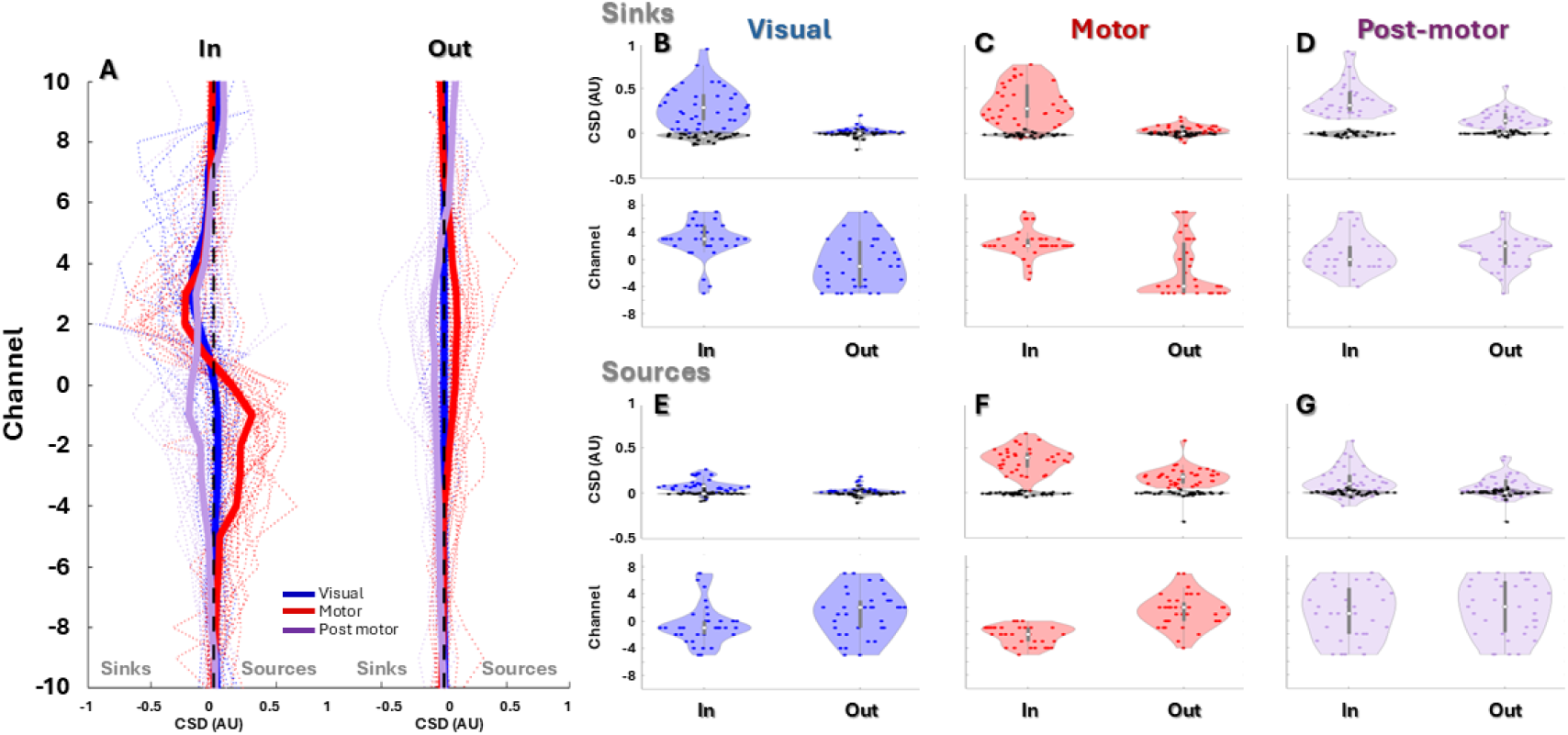
Spatiotemporal CSD profiles in the In and Out task. (A) Median CSD activity across depth during the visual (blue), motor (red), and post-motor (purple) windows, shown separately for targets inside the receptive field (In-RF; left) and outside the receptive field (Out-RF; right). Thick traces represent the median across datasets, and thin traces correspond to individual datasets. (B-G) Violin plots summarizing the distribution of CSD hotspots across datasets. In each panel, the top row shows hotspot magnitude and the bottom row shows hotspot depth for In-RF (left) and Out-RF (right) conditions. Panels B-D show sinks during the visual, motor, and post-motor windows, respectively, whereas panels E-G show sources during the corresponding windows. Colors denote the analysis window (visual: blue; motor: red; post-motor: purple), and black indicates depth-matched baseline values sampled at the corresponding hotspot depth. Baseline hotspot depth is not shown because baseline values were sampled at the depth of the corresponding analysis-window hotspot. Sink values were multiplied by −1 so that sinks and sources are plotted on a common positive scale.

To appreciate these effects in more detail, we show violin plots of the magnitude of sink and source hotspots (Figure 3B-G, top row) and the depth at which they occur (bottom row) across sessions. In these comparisons, blue, red, and purple correspond to the visual, motor, and post-motor epochs, respectively, whereas black denotes the corresponding baseline period for each epoch. The sink values were multiplied by −1 so both sinks and sources are plotted on a common positive scale. During the visual and motor epochs, sink values were higher for in-RF targets than for out-RF targets (visual: 0.30 ± 0.14 AU median ± MAD vs 0.02 ± 0.01 AU, Wilcoxon signed-rank tests p < 0.001; motor: 0.27 ± 0.15 vs 0.04 ± 0.03 AU, p < 0.001). In both epochs, sink values were higher than baseline for in-RF targets (visual baseline: −0.04 ± 0.03 AU, p < 0.001; motor baseline: −0.01 ± 0.02 AU, p < 0.001). Although out-RF values also differed significantly from baseline (visual baseline: −0.00 ± 0.01 AU, p < 0.001; motor baseline: −0.00 ± 0.01 AU, p < 0.001), these responses remained much closer to baseline than those observed for in-RF targets. During the post-motor epoch, sink values remained higher for in-RF than out-RF targets (0.31 ± 0.08 vs 0.14 ± 0.05 AU; p < 0.001) and were higher than baseline in both conditions (in-RF baseline: −0.01 ± 0.01 AU, p < 0.001; out-RF baseline: −0.00 ± 0.01 AU, p < 0.001; Wilcoxon signed-rank tests).

For in-RF targets, sink hotspots occurred at a more superficial depth during the visual epoch (3 ± 1 channels, median ± MAD) than during the motor epoch (2 ± 1 channels), and this difference reached statistical significance (Wilcoxon signed-rank test, p <0.05). Sink hotspots during both the visual and motor epochs were also more superficial than during the post-motor epoch (0 ± 2 channels; visual vs post-motor: p < 0.001; motor vs post-motor: p < 0.05, Wilcoxon signed-rank tests).

During the visual epoch, source modulations were small but higher for in-RF targets (0.07 ± 0.03 AU, median ± MAD) than for baseline (−0.01 ± 0.01 AU; p < 0.001). Although out-RF modulations also differed significantly from baseline (0.02 ± 0.01 AU vs. −0.00 ± 0.01 AU; p < 0.001), they remained much closer to baseline than the in-RF responses.

During the motor epoch, source modulations were larger for in-RF than out-RF targets (0.40 ± 0.08 vs 0.18 ± 0.06 AU, p < 0.001), whereas during the post-motor epoch, source modulations did not significantly differ between in-RF and out-RF targets (0.10 ± 0.08 vs 0.06 ± 0.05 AU, p-value = 0.06; Wilcoxon signed-rank tests). Source modulations exceeded baseline in both conditions (motor: in-RF −0.01 ± 0.01 and out-RF 0.00 ± 0.01 AU, p < 0.001 and p < 0.001, respectively; post-motor: in-RF 0.00 ± 0.01 and out-RF 0.00 ± 0.01 AU, p < 0.001 and p < 0.001, respectively; Wilcoxon signed-rank tests). Importantly, source amplitudes were lower in the post-motor than in the motor epoch in both conditions (in-RF: p < 0.001; out-RF: p < 0.001; Wilcoxon signed-rank tests).

During the motor epoch, source hotspots for in-RF targets occurred at deeper depths than those for out-RF targets (−2 ± 1 vs 2 ± 2 channels; Wilcoxon signed-rank test, p <0.001) and were also deeper than the corresponding in-RF sink hotspots ((−2 ± 1 vs 2 ± 2 channels; p <0.001).

Overall, CSD modulations showed distinct laminar and spatial profiles across epochs. During the visual epoch, sink activity was restricted to in-RF targets and confined to superficial layers. During the motor epoch, sink activity remained restricted to in-RF locations, whereas source activity was observed for both in-RF and out-RF targets, with sources occurring at deeper depths than sinks. During the post-motor epoch, sink activity was present for both in-RF and out-RF conditions. Thus, visual responses were highly localized both spatially and across SC depth, whereas motor-related activity recruited both superficial and deeper layers and engaged activity outside the in-RF location. Negative deflections (sinks) are generally interpreted as reflecting local excitation, whereas positive deflections (sources) are commonly associated with return currents or inhibition. Accordingly, these patterns are consistent with localized excitation during the visual epoch, followed by broader circuit recruitment during the motor and post-motor periods.

### Spatial extent of CSD activity differs across visuomotor epochs

The analyses above interpret excitation and inhibition signatures in the CSD for two extremes target conditions (in-RF and out-RF), leaving unresolved how these activity patterns vary across intermediate target locations. We therefore next examined CSD activity during the 8-target task, which samples locations spanning the intervening spatial range. Figure 4 shows, for an example session, the trial-averaged CSD activity across depth for the eight target locations during the visual (left column) and motor (right column) epochs. The panels are arranged according to target position, preserving the spatial layout of the target array. The corresponding LFP and spiking activity for this example session are shown for the visual and the motor epochs in Supplementary Figures 1 and 2, respectively.

**Figure 4.**
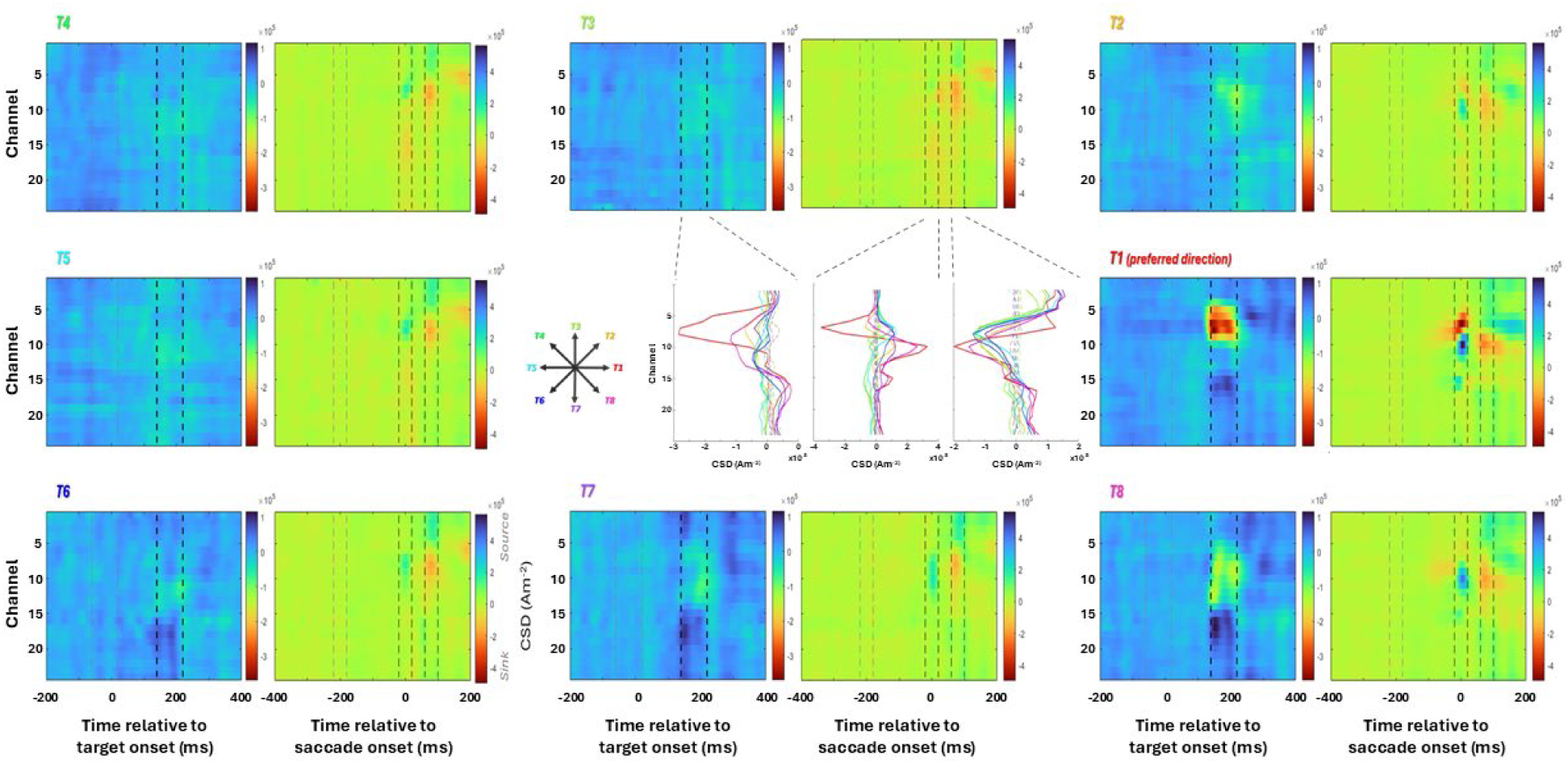
Spatiotemporal CSD profiles in the 8-target task from an example session. CSD activity across depth aligned to target onset (left) and saccade onset (right) is shown for each of the eight target locations (T1-T8). T1 corresponds to the preferred direction and is located within the response field (RF). Red colors indicate sinks and blue colors indicate sources. Vertical gray and black dashed lines indicate the baseline and analysis windows, respectively. The middle panels show average CSD activity across depth during the visual (left), motor (middle), and post-motor (right) windows (solid lines), together with the corresponding baseline activity (dashed lines). Colors correspond to target locations T1-T8, as labeled in the target-location schematic.

During the visual epoch, a prominent sink modulation (red hotspot) was observed in superficial layers when the target was presented in RF (i.e., “preferred direction”), accompanied by a source (blue hotspot) at deeper depths. Weaker sink activity was observed for targets adjacent to the RF (see Targets 2 and 8), whereas little or no modulation was present for more distant locations. Thus, visual-epoch CSD activity was spatially restricted, with modulation strongest in or near the RF and decreasing with increasing distance from the preferred location, as summarized in the middle-left panel, which shows the average CSD activity across depth during the visual window for all target locations. During the motor epoch, a sink modulation (red hotspot) paired with a deeper source (blue hotspot) was again observed when the saccade was directed toward the RF. In contrast to the visual epoch, however, the source activity was present across a broader range of target locations, although with larger amplitudes for movements toward the RF. This indicates that motor-related CSD activity was distributed across a wider portion of the SC map and across a broader range of laminar depths than visual responses, as summarized by the average CSD across depth (middle panel). In contrast to sink modulation during the visual and motor epochs, sink activity during the post-motor period (red hotspot) was not restricted to movements directed toward the RF but was present for all saccade directions, with relatively similar laminar patterns across targets. This indicates that post-motor CSD activity was broadly distributed across the SC map. This pattern is summarized by the average CSD across depth during the post-motor window for all target locations (middle panel). LFP signals showed patterns consistent with these observations, with visual responses largely restricted to the RF and motor-related activity present across a broader range of target locations (see Supplementary Figures 1 and 2 for spike density and LFP signals from the same example session). In contrast, spiking activity remained largely confined to the RF in both epochs, providing limited information about activity at other locations (also shown in Supplementary Figures 1 and 2).

Across datasets, CSD modulations were consistent with the patterns observed in the example session. Median profiles across depth (Figure 5, thick traces) revealed that sink activity during the visual and motor epochs remained strongest for targets in the response field, whereas post-motor sink activity was more broadly distributed across target locations. A similar pattern was observed for sources: motor-epoch sources were strongly tuned toward the response field, while sources during the visual and post-motor epochs showed weaker spatial selectivity. This difference in target selectivity is summarized in the polar plots (Figure 5), which show that sink modulations during the visual and motor epochs, and source modulations during the motor epoch, were strongly biased toward the RF (T1), whereas sink and source modulations during the post-motor epoch were more broadly distributed across directions.

**Figure 5.**
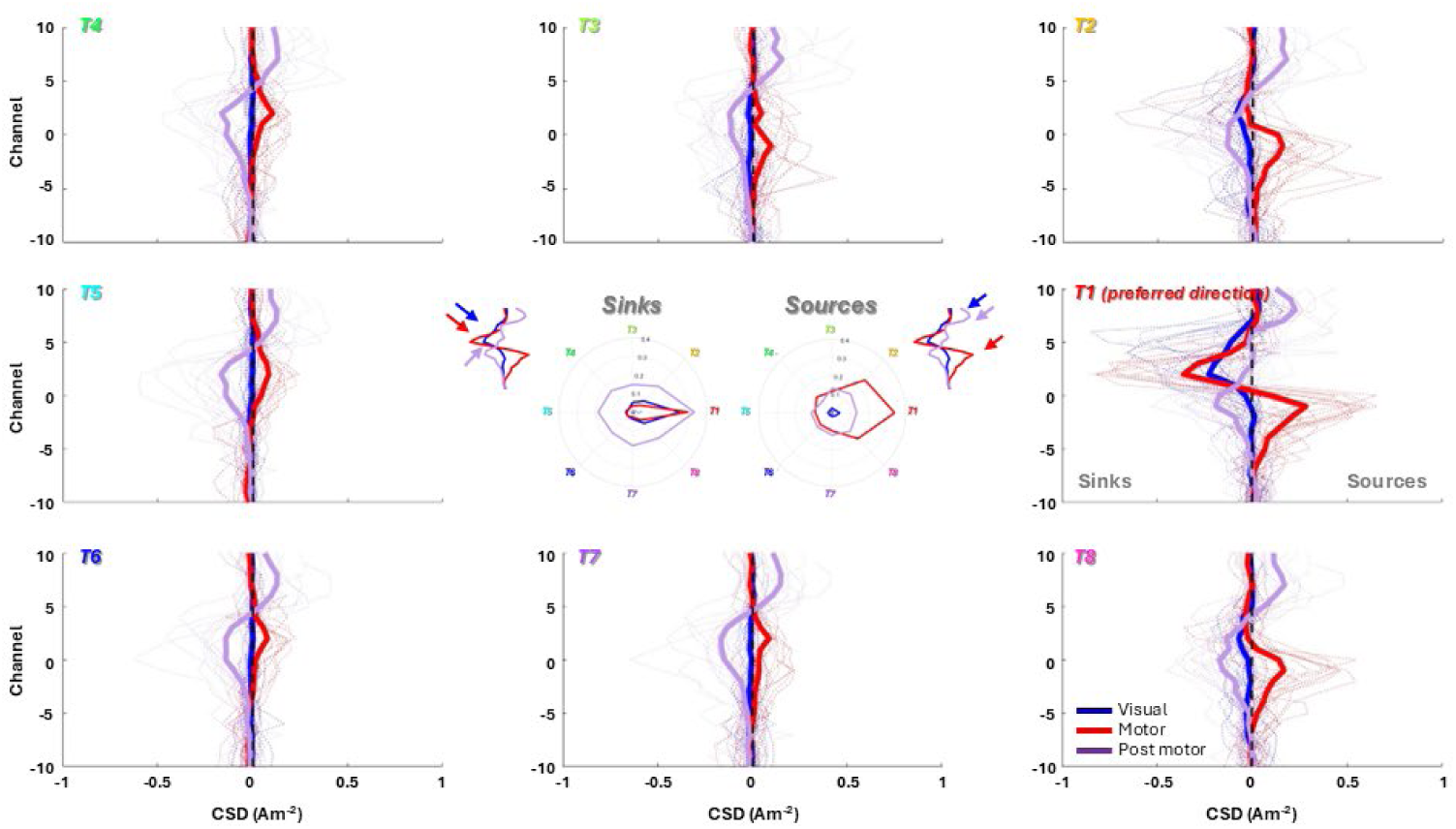
Spatiotemporal CSD profiles in the 8-target task across datasets. Median normalized CSD activity across depth is shown for each of the eight target directions during the visual (blue), motor (red), and post-motor (purple) windows. Thick traces represent the median across datasets, and thin traces correspond to individual datasets. In the depth profiles, negative values correspond to sinks and positive values to sources. Polar plots summarize the spatial tuning of CSD activity across target directions. Colored traces correspond to CSD values during the visual (blue), motor (red), and post-motor (purple) windows. For polar plots, sink values were multiplied by −1 so that sinks and sources are displayed on a common positive scale for visualization.

Figure 6A shows the CSD values (upper row) and the channels at sink hotspots (bottom row) for the different target locations during the visual (left, blue), motor (middle, red), and post-motor (right, purple) responses across sessions. For all epochs, CSD values were significantly higher when the target was in the RF (visual: 0.52 ± 0.21 AU; motor: 0.30 ± 0.26 AU; post-motor: 0.33 ± 0.12 AU, median ± MAD) compared with their respective baseline (visual: −0.05 ± 0.03 AU; motor: 0.00 ± 0.02 AU; post-motor: 0.00 ± 0.02 AU, Wilcoxon signed-rank tests, p < 0.001, p < 0.001, and p < 0.001, respectively) and decreased with increasing distance from the preferred location (target opposite the RF (i.e., target 5, 180° apart): visual: 0.04 ± 0.01 AU; motor: 0.03 ± 0.02 AU; post-motor: 0.19 ± 0.12 AU; Wilcoxon signed-rank tests, p < 0.001, p < 0.001, and p < 0.05, respectively). Significant correlations were found between sink modulations and target location during the visual (circular-linear correlation coefficient = 0.41, p < 0.001), motor (circular-linear correlation coefficient = 0.34, p < 0.001), and post-motor epochs (circular-linear correlation coefficient = 0.20, p < 0.05).

**Figure 6.**
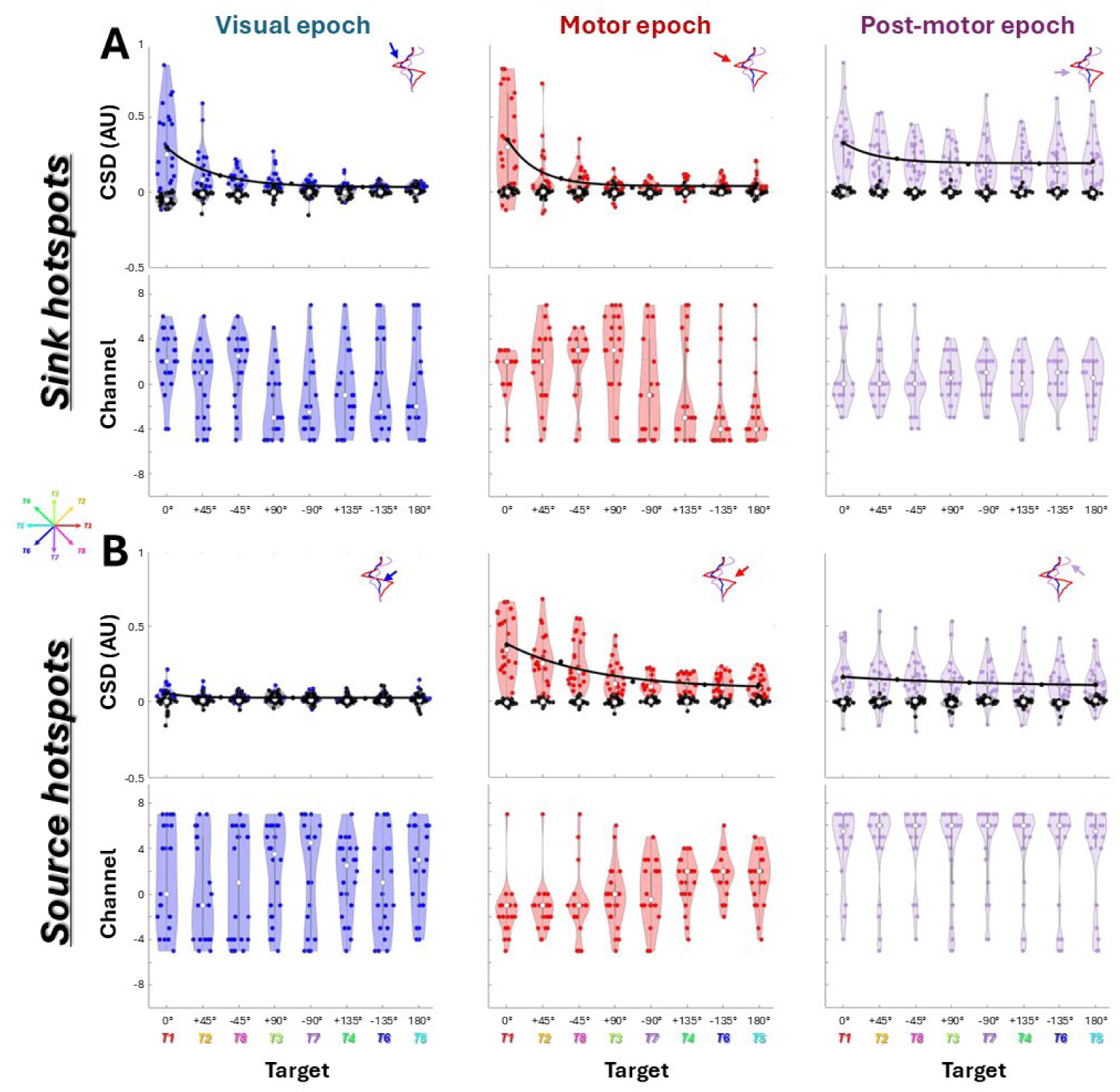
Sink and source hotspots in the 8-target task across datasets. (A) Sink and (B) source hotspots are shown across target locations during the visual (left, blue), motor (middle, red), and post-motor (right, purple) epochs. For each epoch, the top row shows hotspot magnitude, and the bottom row shows hotspot depth (channel) across datasets. Targets are ordered by angular distance from the preferred direction (T1). Black points indicate depth-matched baseline CSD values. Baseline hotspot depth is not shown because it is defined to match the hotspot depth. Black lines show exponential fits of hotspot magnitude as a function of angular distance from the preferred direction, computed from the median values across datasets at each target location. Decay constants (κ) were estimated independently for each dataset and used for statistical comparisons.

To further characterize the spatial extent of sink hotspots, we quantified the decay constant *k* of the CSD modulation across target locations. Larger *k* values indicate a steeper spatial decay and a more spatially restricted profile, whereas smaller *k* values indicate a shallower decay and a broader spatial profile. Decay constants were significantly greater than zero during the visual, motor, and post-motor epochs (Wilcoxon signed-rank tests, all *p* < 0.001; see supplementary figure 3), indicating that sink magnitude decreased with increasing distance from the RF across all three epochs. Decay was significantly higher during the motor than during the visual epoch (*p* < 0.01; Wilcoxon signed-rank test), indicating a more spatially restricted profile during the motor epoch (see the sink polar plot in Figure 5, where the motor (red) profile shows sharper spatial tuning than the visual (blue) profile). Decay did not differ significantly between the motor and post-motor (p-value = 0.12) or visual and post-motor epochs (p-value = 0.31; Wilcoxon signed-rank tests).

Despite similar median depths during the visual and motor epochs (visual: 2 ± 2 channels; motor: 2 ± 1 channels; median ± MAD), visual sink hotspots occurred significantly more superficially than motor sink hotspots (Wilcoxon signed-rank test, p<0.01), with mean depths of 2.00 and 1.06 channels, respectively.

Figure 6B shows the CSD values (upper row) and the channels at source hotspots (bottom row) for the different target locations for the visual (left, blue), motor (middle, red), and post-motor (right, purple) responses across sessions. During the visual epoch, source values were higher when the target was in the RF (target 1: 0.04 ± 0.02 AU, median ± MAD) and decreased with target eccentricity, reaching 0.02 ± 0.01 AU at the opposite target location (target 5, 180° apart; Wilcoxon signed-rank test, p < 0.01), where values were similar to baseline (0.02 ± 0.01 AU).

Moreover, a significant correlation was found between the source modulations and the target location (circular-linear correlation coefficient = 0.21, p< 0.05). When the target is presented in the RF, the source hotspots appear either in more superficial (10 sessions, median depth= 6 ± 1) or deeper layers (12 sessions, median depth= −2 ± 1.5) (see bimodal distribution) than sink hotspots during the visual epoch (median depth= 2 ± 2 in fig. 6A).

During the motor and the post-motor epochs, the source values are higher when the target is in the RF (0.34 ± 0.17 AU median ± MAD and 0.13 ± 0.08 AU for motor and post-motor epoch, respectively) and decrease with RF proximity (0.09 ± 0.06 AU and 0.11 ± 0.07 AU, for target located 180° apart (i.e., target 5), Wilcoxon signed-rank test, p < 0.001 and p < 0.01) but stay always higher than baseline (motor= −0.01 ± 0.01 AU and post-motor= −0.00 ± 0.01 AU for target 1; motor= −0.00 ± 0.01 AU and post-motor= 0.01 ± 0.01 AU for target 5) during the motor and post-motor epoch, respectively). CSD values are different from baseline values (black) for all saccade direction (p<0.05, Wilcoxon signed rank test) during both the motor and the post-motor epoch. Significant correlations were found between the source modulations and the target location for the motor epoch (circular-linear correlation coefficient = 0.49, p < 0.001) but not for the post-motor epochs (circular-linear correlation coefficient = 0.12, p-value = 0.30). Note that the later correlation is weaker.

We next quantified the spatial extent of source hotspots using the same decay analysis across target locations. As for sinks, larger κ values indicate a steeper spatial decay and a more spatially restricted profile. Decay constants were significantly greater than zero during the visual, motor, and post-motor epochs (Wilcoxon signed-rank tests, all p < 0.001; see supplementary figure 4), indicating that source magnitude decreased with increasing distance from the RF across all three epochs. Decay did not differ significantly between the visual and motor (Wilcoxon signed-rank tests; p-value = 0.33), motor and post-motor (p-value = 0.28), or visual and post-motor epochs (p-value = 0.91).

Interestingly, During the motor epoch, when the target is presented in the RF, the source hotspots appear in deeper layers (median depth= −1 ± 1 MAD) than the sink hotspots (median depth= 2 ± 1 MAD) and were also deeper than the source hotspots observed for the target outside the RF (median depth = 2 ± 1.5 MAD for Target 8). During the post-motor epoch, the source hotspots appear approximatively at the same layers for all saccade directions (median depth= 5.5 ± 1.5 MAD and 5.5 ± 1.5 MAD for target 1 and 5, respectively).

Overall, the 8-target task showed that visual responses were spatially restricted and confined to superficial layers, whereas motor and post-motor activity was distributed across a broader range of target locations and depths.

## Discussion

Using laminar recordings and CSD analysis in the SC of behaving non-human primates, we uncovered a state- and depth-dependent reconfiguration of neural activity during visuomotor transformation. Figure 7 summarizes how these distinct regimes of excitation and inhibition evolve across behavioral states and SC depth. During sensory encoding (Fig. 7, left panels), activity is dominated by spatially confined excitation in superficial layers, supported by a focal sink observed only when the target fell within the receptive field of the recorded SC site. During movement execution (Fig. 7, middle panels), the circuit shifts to a configuration characterized by tightly coupled sinks and deeper sources for preferred saccades, alongside widespread source activity for non-preferred directions, consistent with local excitation and distal inhibition. Following movement completion (Fig. 7, right panels), the network in the intermediate layers enters a direction-invariant state marked by distributed sink activity, consistent with widespread reorganization of collicular excitability. This global excitation is accompanied by direction-invariant inhibition in the superficial layers. These findings demonstrate that excitation and inhibition in the SC do not operate as a fixed center-surround architecture. Instead, the circuit transitions between temporally defined functional states that reorganize patterns of excitation and inhibition as sensorimotor transformation transpires. By resolving transmembrane currents across SC layers, our results provide population-level evidence that sensorimotor transformation is supported by a dynamic reconfiguration of excitation and inhibition across space and time.

**Figure 7.**
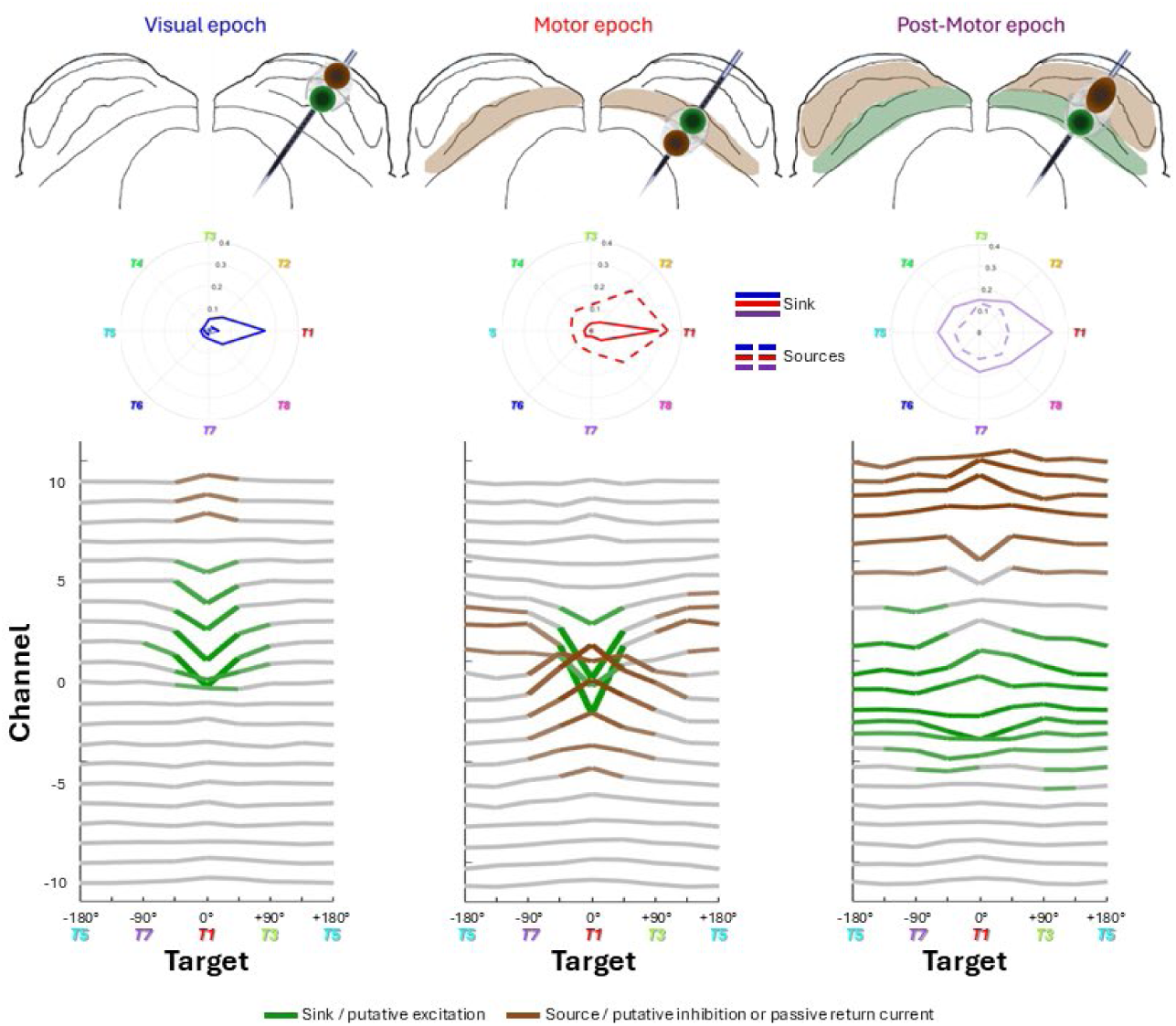
Schematic summary of the spatial organization of excitation and inhibition across the SC during different behavioral epochs. Columns correspond to the visual (left), motor (middle), and post-motor (right) epochs. Top: Schematics illustrate the proposed spatial organization of activity across the SC map. Green and maroon circles indicate the locations of the strongest sink and source hotspots, respectively. Shaded regions illustrate the inferred spatial extent of the underlying processes, with green representing putative excitation and maroon representing putative inhibition or passive return currents associated with a dipole. Middle: Polar plots summarize the spatial distribution of sink (solid lines) and source (dashed lines) magnitudes across the eight target directions during the visual (blue), motor (red), and post-motor (purple) epochs; thinner lines indicate the corresponding depth-matched baseline CSD activity. Sink values were multiplied by −1 so that sinks and sources are displayed on the same positive scale for visualization and comparison. Bottom: Median CSD profiles across depth are shown as a function of angular distance from the preferred direction (T1). Green traces indicate sinks, maroon traces indicate sources, and gray traces indicate CSD profiles without prominent sink or source activity. During the visual epoch, activity is spatially restricted to the RF location and dominated by a superficial sink, with a source observed in some datasets that may represent the return current of a local dipole. During the motor epoch, a sink and source are present at the RF location, whereas sources extend across non-RF locations without corresponding sinks, consistent with spatially broader inhibition. During the post-motor epoch, sinks and sources extend across target locations, consistent with widespread, layer-specific excitation and inhibition.

### CSD modulation after stimulus presentation

During the visual epoch, we consistently observed a current sink in the superficial layers of the superior colliculus, flanked by upward and/or downward current sources, forming a vertically organized sink-source pattern (see Figs. 2E and 4). This configuration emerged shortly after target onset and was observed only when the target appeared within the RF of the recorded SC site. The laminar confinement of the sink to superficial layers is consistent with excitatory inputs arising from the retina and visual cortical areas, which are known to terminate predominantly within these layers. Thus, this sink likely reflects retinotopically specific excitatory drive entering the superficial SC. The absence of comparable modulation for targets outside the receptive field further indicates that visual excitation remains spatially constrained during the sensory epoch.

The spatial confinement of visual activity was somewhat unexpected given previous findings from multi-target paradigms. When a visual stimulus is presented within the RF of an SC neuron, its visually evoked spiking response is reduced by the simultaneous presentation of additional stimuli outside the RF, an effect commonly attributed to lateral inhibitory interactions across the collicular map (Basso & Wurtz 1998, McPeek & Keller 2002). Such findings suggest that activation of distant SC representations can suppress activity at the RF location during sensory processing. However, this interpretation has been based primarily on changes in spiking activity and does not establish whether the suppression arises from inhibitory interactions within the SC itself or from reduced excitatory drive arriving from upstream structures. Reduced visual responses in the presence of multiple stimuli have been observed across several oculomotor structures. In FEF and LIP, the initial visual response is reduced when multiple stimuli are presented compared with a single stimulus (Schall et al 1995, Thomas & Paré 2007). Similarly, in the substantia nigra pars reticulata, the visually evoked pause decreases as the number of potential targets increases and can even be absent (Basso & Wurtz 2002). In our data, presentation of a single target outside the RF produced no detectable modulation of the LFP or CSD at the recorded RF location, providing no evidence for distal inhibition under these conditions. Importantly, however, our paradigm activated only one target representation at a time. We therefore cannot exclude the possibility that simultaneous activation of multiple locations recruits lateral inhibitory interactions that are absent when a single distant location is activated. Direct comparison of single- and multi-target conditions using laminar CSD recordings will be necessary to determine whether multi-target suppression reflects inhibition generated within the SC or changes in the excitatory inputs reaching it.

The spatial constraints of visual activity may also reflect the cellular organization of the superficial SC. Neurons in these layers include narrow- and wide-field vertical cells, whose dendritic arbors can sample visual inputs over different spatial extents while their output is organized primarily toward local and deeper collicular layers (Basso & May 2017, Isa et al 1998, Laemle 1981, Langer & Lund 1974). Despite differences in the spatial extent of their dendritic arbors, the predominantly local and interlaminar organization of their outputs may favor transmission between collicular layers rather than broad lateral interactions across the collicular map, potentially contributing to the spatially confined excitation and absence of the distal inhibitory pattern observed during the motor epoch.

The sink in the superficial SC has previously been leveraged as a physiological landmark for laminar alignment across recording sessions, allowing recordings from different SC penetrations to be aligned according to a common reference depth before averaging (Bourrelly et al 2023, Lee et al 2020, Massot et al 2019), but the associated transmembrane current dynamics were not directly examined. Laminar alignment has previously been based on the transition between a superficial sink and a deeper source (2019). Notably, the CSD profiles in that study also exhibited a superficial source above the sink, mirroring the flanking source configuration observed here. (Massot et al., 2019). Notably, the CSD profiles in that study also exhibited a superficial source above the sink, mirroring the flanking source configuration observed here. A comparable sink-source configuration has also been described in the ferret SC following stimulus onset, along with a second, deeper sink-source pair emerging approximately 100 µm below the initial pattern ∼40 ms later (Stitt et al 2013). We observe hints of a similar secondary sink-source configuration in our data (see Fig. 4, T1), although in our recordings these components appear partially merged. This difference may reflect the continuous target presentation used in our task, in contrast to the brief flashes (2ms) employed in their paradigm. The presence of multiple sink-source pairs raises the possibility that distinct excitatory inputs from different sources converge in different SC layers and with different temporal profiles.

In a subset of sessions, we also detected a transient modulation preceding the dominant visual epoch sink in superficial layers (see Fig. 2C, in-RF column). This early component appeared as a brief source in the CSD signal accompanied by a small positive deflection in the LFP. One plausible interpretation is that it reflects return currents or dendritic processing associated with initial synaptic activation before the peak excitatory drive. Transient early components of this kind have been reported in laminar recordings and attributed to the spatial arrangement of synaptic inputs along dendritic arbors in sensory cortex (Kajikawa & Schroeder 2011, Schroeder et al 1998). Although we did not quantify this feature because it was not consistently present across sessions, its occurrence suggests that additional subthreshold processes may precede the principal excitatory response during visual processing.

### CSD modulation during saccade generation

Consistent with the hypothesis that excitation-inhibition balance reorganizes across behavioral stages, CSD patterns during the motor epoch revealed distinct laminar signatures depending on saccade direction. When saccades were directed into the RF, a sink emerged in intermediate layers paired with a deeper source (see Figs 2F and 4). This configuration is consistent with excitatory input to premotor SC neurons accompanied by coordinated inhibitory and/or return-current activity. The laminar position of this motor-related sink aligns with the distribution of buildup and burst neurons in the intermediate layers (Massot et al 2019, Mohler & Wurtz 1976) and with the anatomical organization of motor output pathways projecting to the brainstem saccade generators (Basso & May 2017, Gandhi & Katnani 2011, Wurtz & Albano 1980).

Conversely, when saccades were directed away from the RF, source activity was observed in intermediate layers without a corresponding sink. This polarity reversal is consistent with active inhibitory recruitment rather than simply a reduction in the excitatory drive. Potential sources of this suppression include local GABAergic circuitry, commissural interactions between the two colliculi (Munoz & Istvan 1998, Takahashi et al 2010), or descending influences from cortical and basal ganglia pathways, including the substantia nigra pars reticulata (Basso & Sommer 2011, Hikosaka et al 2000, Hikosaka & Wurtz 1983). These saccade direction-dependent laminar patterns reveal a spatial organization characterized by local excitatory recruitment at the selected movement site and suppression of neural activity across competing regions of the collicular map, providing a potential mechanism for competitive selection in which the selected motor command is amplified while competing movement vectors are suppressed, consistent with previous models of collicular competition (Dorris et al 2007, McPeek & Keller 2002, Munoz & Istvan 1998). Importantly, this motor-related inhibitory signature was evident even in the absence of spiking activity (see supplementary figure 2), indicating that suppression during saccade generation involves structured subthreshold processes that are not captured by firing-rate analyses.

### CSD modulation after saccade generation

The post-motor period exhibited a laminar CSD pattern that was distinct from both the visual and motor epochs (see Figs 2E-F and 4). Rather than returning to baseline following saccade execution, the SC displayed a robust dipole configuration characterized by a sink in intermediate layers paired with a superficial source, that was observed across all saccade directions. The directional invariance of this modulation suggests that it reflects a global circuit process rather than vector-specific planning or execution. One possibility is that this activity reflects rebound excitation or circuit disinhibition following widespread inhibitory engagement during saccade execution. Rebound-like phenomena have been described in sensorimotor circuits and may contribute to resetting network excitability after action (Hikosaka et al 2000, Jagadisan & Gandhi 2017b). Alternatively, the post-movement signal may reflect corollary discharge or feedback from oculomotor and cortical pathways involved in monitoring movement outcomes and updating internal state representations (Sommer & Wurtz 2001, Wurtz et al 2011). Regardless of its precise origin, the temporally structured and spatially widespread nature of this post-motor activity demonstrates that the SC remains actively engaged after movement execution, supporting the view that sensorimotor transformation extends beyond saccade initiation and includes processes involved in resetting or updating the network for subsequent behavior.

### Dynamic reorganization of excitation-inhibition balance

Our results refine and extend previous models of excitation-inhibition organization in the SC by showing that these interactions are not fixed architectural motifs but dynamically reconfigured across behavioral epochs. Slice studies have revealed structured spatial motifs, including center-surround organization in superficial layers and broader excitatory-inhibitory interactions in deeper layers (Isa & Hall 2009, Meredith & Ramoa 1998, Phongphanphanee et al 2008, Phongphanphanee et al 2014). These studies suggest that visual input engages local excitation accompanied by distal inhibition. However, we did not observe comparable inhibitory signatures during the visual epoch outside the receptive field. Instead, visual responses were dominated by spatially localized excitation with minimal modulation at distant sites. This discrepancy may reflect differences between intact behaving networks and reduced slice preparations, including ongoing cortical and subcortical inputs, behavioral state, and the continuous visual stimulation used here.

In contrast, during the motor epoch, our findings converge with previous reports of broader collicular interactions. In vivo studies in cats and monkeys have shown lateral inhibitory interactions and competitive dynamics across the SC map (Munoz & Istvan 1998, Takahashi et al 2010), consistent with models of vector competition (Hoffmann & Straschill 1971, Mascetti & Arriagada 1981). Here, we directly visualize this competition through the laminar organization of population activity: preferred saccades were associated with focal excitation coupled with deeper inhibitory or return-current activity, whereas nonpreferred directions exhibited isolated source signatures consistent with active suppression. Thus, whereas slice studies reveal the potential circuitry underlying center-surround organization, our data demonstrate how this circuitry is differentially recruited as behavior unfolds.

Importantly, most previous in vivo work relied on anesthesia, pharmacological inactivation, or transient perturbations (Dorris et al 2007, Haider et al 2013, Jagadisan & Gandhi 2017a, Munoz & Istvan 1998, Populin 2005, Skoblenick & Everling 2012), which can alter excitation-inhibition balance and obscure state-dependent dynamics. By contrast, our laminar CSD recordings in behaving primates reveal that excitation dominates during sensory encoding, whereas inhibitory and disinhibitory processes become prominent during motor selection and post-movement phases. These findings suggest that excitation-inhibition balance in the SC is not a static center-surround mechanism continuously active across epochs, but rather a behaviorally gated process that reorganizes as the circuit transitions from stimulus encoding to action selection and network resetting.

### LFP, CSD, and dipole considerations

Because extracellular potentials spread through conductive tissue (Buzsaki et al 2012, Mitzdorf 1985), interpretation of LFP signals alone does not reveal the spatial origin of the underlying current generators. Current source density analysis, by computing the second spatial derivative of extracellular potentials (Freeman & Nicholson 1975, Nicholson & Freeman 1975, Pettersen et al 2006), allows laminar resolution of net transmembrane currents. Although sinks and sources are often interpreted as excitation and inhibition, respectively, such interpretations must use caution because return currents can produce similar signatures (Buzsaki et al 2012, Kajikawa & Schroeder 2011). In our data, some coupled sink-source patterns likely reflect such dipole organization (Figure 5, motor epoch; red trace in the middle-right panel), where the RF condition exhibits a sink accompanied by a deeper source. However, the presence of unpaired source activity without a corresponding sink is consistent with active inhibitory processes contributing substantially to the observed laminar profiles. Accordingly, the laminar patterns reported here are best interpreted as reflecting the combined influence of passive current spread and active membrane processes whose relative contributions vary across behavioral epochs.

In summary, our results provide a depth-resolved view of population activity in the superior colliculus as sensory information is transformed into motor commands. Visual processing is dominated by spatially localized excitation, motor execution recruits tightly coordinated excitatory-inhibitory interactions, and movement completion engages widespread activity consistent with network resetting or state updating. These observations suggest that the computational roles of excitation and inhibition evolve across the sensorimotor transformation: early visual processing favors faithful representation of sensory input through localized excitation with little evidence of widespread suppression, whereas motor selection recruits inhibitory interactions to resolve competition and stabilize the selected movement. Many of these dynamics unfold below spike threshold, revealing that a substantial portion of sensorimotor computation is hidden from conventional measures of neuronal output. Together, these findings demonstrate that the SC is not governed by a static circuit architecture, but by a dynamically reconfigured network in which patterns of excitation and inhibition are continuously reshaped to guide behavior.

## Methods

### Animal preparation

Four adult male rhesus macaques (Macaca mulatta; monkeys BB, BL, SU, and DU) participated in this study. A stainless-steel recording chamber (Narishige) angled 40° posterior with respect to vertical was implanted on each animal during a sterile surgery. The details of the surgical procedure have been described previously (Katnani & Gandhi 2012, Massot et al 2019). Electrode insertions through this chamber are oriented perpendicular to the SC surface and follow the dorsoventral axis, passing through regions where neurons share similar response fields. All experimental procedures were approved by the University of Pittsburgh Institutional Animal Care and Use Committee.

### Behavioral paradigm and data analysis

Stimulus display, behavioral paradigms, and all data acquisition were managed with a real-time LabView-based controller interface (Bryant & Gandhi 2005) from monkey BB, BL and SU and a Psychtoolbox-based interface for monkey DU. An LED-backlit flat screen television (120 frames/s, 1080p resolution) was used to present white targets (4 x 4 pixels) on a dark gray background. The stimulus subtended ∼0.5° of visual angle. To synchronize timing across experimental stages, a white square was simultaneously flashed at the screen corner and recorded via a photodiode. This region of the monitor was covered so that the white square and photodiode were not visible to the animal. Eye position was sampled at 1 kHz with an infrared eye tracker (EyeLink 1000, SR Research) calibrated at the beginning of each session by having the animal fixate targets presented at known locations on the monitor. The head was restrained by using a thermoplastic mask (Drucker et al 2015).

During each recording session, animals performed a center-out, visually guided delayed saccade task. Each trial began with fixation of a central point. After a variable delay period (600-1500 ms), a peripheral target appeared while fixation was maintained. Following a go cue (fixation point offset), animals were required to execute a saccade to the target location and received a liquid reward for landing within an acceptance window centered on the peripheral target, typically 2°-6° in radius but adjusted when necessary based on calibration quality and target eccentricity. Only successful trials were included in the analysis. Two variants of this delayed saccade task were performed on separate recording days. In the In-and-Out task, the peripheral target was presented either within the neuron’s response field (In-RF) or at the diametrically opposite location (Out-RF). Targets within the RF were presented more frequently (80%) than Out-RF targets (20%). This task was performed in BB, BL and SU (data were previously reported in part (Bourrelly et al 2023). In the 8-targets task, targets were presented at eight locations equally spaced at 45° intervals along an imaginary circle at constant eccentricity. One of these locations was designated Target 1, and the remaining targets were numbered sequentially in a counterclockwise direction. For Monkeys SU and DU, Target 1 corresponded to the RF location. For Monkey BL, targets were arranged at a fixed eccentricity of 10°, positioned along the cardinal and oblique directions, and the location closest to the estimated RF was defined as Target 1. All target locations were presented with equal probability. A subset of these was previously reported in (Heusser et al 2022).

All trials were analyzed offline using custom code written in MATLAB (Mathworks, Inc). The onset and offset of the horizontal and vertical components of each saccade were detected using a velocity threshold of 30°/s. Each trial was also visually inspected, and the detected onset and offset were manually adjusted when necessary.

### Neurophysiological recordings and microstimulation

Neural activity along the dorsoventral extent of the SC was recorded using laminar electrodes. A hydraulic microdrive (Narishige) was used to advance either a 16-channel linear probe (Alpha-Omega; 150 μm intercontact spacing; ∼300 μm shaft diameter; ∼1 MΩ impedance per contact) in Monkeys BB and BL, or a 24-channel linear probe (S-probe, Plexon; 150 or 200 μm intercontact spacing; ∼210 μm diameter) in Monkeys SU and DU. After the probe reached the intermediate layers of the SC, its position was adjusted to maximize the number of channels exhibiting clear multiunit activity. To estimate the preferred movement field at each penetration, electrical microstimulation was applied on selected contacts (40 μA, 400 Hz, 200 ms; biphasic pulses, 200 μs phase duration, 17 μs interphase interval), and the evoked eye movements were averaged. The resulting mean saccade vector (direction and amplitude) was taken as an estimate of the preferred response field for neurons recorded across the probe contacts. At the conclusion of each recording session, the stimulation protocol was repeated systematically across all channels while behavior was monitored (see also (Katnani & Gandhi 2012, Massot et al 2019)).

### Neural data collection and processing

Neurophysiological signals were acquired using a Scout recording system (Ripple). Neural recordings were synchronized at the start of each trial with our behavioral control systems. Raw signals from each channel were processed offline to separate local field potentials (LFPs) from high-frequency activity. LFP signals were obtained using a low-pass filter at 250 Hz, while higher-frequency components were extracted using a high-pass filter at 250 Hz. Spike times were identified by applying a standard threshold-crossing procedure. To obtain continuous spike-density waveforms (see examples in Fig. 2A and B), spike trains were convolved with a kernel approximating an excitatory postsynaptic potential, using growth and decay time constants of 1 and 20 ms, respectively (Thompson et al 1996). LFP signals (see examples in Fig. 2C and D) were further bandpass filtered between 0.7 and 30 Hz (Bourrelly et al 2023). In total, 42 and 29 recording sessions were obtained for the In-and-Out task and the 8-target task, respectively. Sessions were excluded when LFP signals were excessively noisy, the signal-to-noise ratio was insufficient across recording contacts, or an insufficient number of trials was available for a given target condition. After exclusion, we report analyses from 35 and 22 sessions across the two tasks.

Current source density (CSD), the second spatial derivative of the LFP, was computed from trial-averaged LFP signals separately for the visual (–400 to 600 ms relative to target onset) and motor (–600 to 400 ms relative to saccade onset) epochs. The iCSD method implemented in the CSDplotter toolbox (https://github.com/espenhgn/CSDplotter) was used to estimate the CSD signal. Analyses were then restricted to three time-windows of interest: the visual epoch (140-220 after target onset for Monkeys BB, BL, and SU, and 40-120ms for Monkey DU), the motor epoch (20 ms before to 20 ms after saccade onset), and the post-motor epoch (60-100 ms after saccade onset). Data from Monkey DU were corrected using the photodiode signal, whereas photodiode recordings were unavailable for Monkeys BB, BL, and SU. For these three monkeys, the stimulus appeared on the monitor approximately 100 ms after the recorded target-onset time, accounting for the 100-ms difference in the visual analysis window. Baseline periods for the visual and motor analyses were defined as activity occurring from 60 ms before to 20 ms after target onset, and from 220 to 180 ms before saccade onset, respectively. Full-epoch CSD plots were displayed in absolute units for visualization purposes. However, prior to averaging within the defined analysis windows and before any quantitative analyses across conditions or sessions, CSD values were normalized. Normalization was performed separately for each recording session by dividing all visual- and motor-aligned CSD values across the different target conditions by the maximum absolute CSD value observed across both alignments and all target locations within that session. All statistical analyses were performed on these normalized, window-averaged CSD signals.

### Depth alignment of multiple sessions

Our analyses focus on neural activities across SC layers (depth) within a session and then averaging the data across sessions. This requires a procedure to spatially “shift” datasets so that all sessions are referenced to the same SC layer. We used an objective method based on the similarity of features in the current source density (CSD) representation during the motor epoch, as described previously (Bourrelly et al 2023). Specifically, alignment was performed by identifying the depth corresponding to the consistent sink-source transition observed around saccade onset in the motor epoch CSD profile. The channel closest to this reversal was taken as the reference depth, and all channels within a session were expressed relative to this reference before averaging across sessions.

### Spatial decay and statistical analysis

To quantify the spatial decay of sink and sources magnitude across target directions, responses were first summarized per session using the average magnitude across trials for each of the eight target locations. To reduce directional redundancy, responses at ±45°, ±90°, and ±135° were averaged, yielding five angular distances from the preferred direction (0°, 45°, 90°, 135°, and 180°). For each session and condition, decay was modeled using an exponential function:

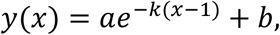

where *x* denotes the 5 target positions along the ordered axis (0° to 180°), *a* is amplitude, *b* is baseline, and *k* is the decay constant. Parameters were estimated independently for each session using nonlinear least-squares optimization with positivity constraints on *a* and *k*.

The decay constant *k* was used as a session-wise measure of spatial sharpness, with larger values indicating more abrupt decay (i.e., sharper spatial tuning) and smaller values indicating shallower decay (i.e., broader spatial tuning). Decay constants were estimated independently for each recording session. Wilcoxon signed-rank tests were used to determine whether decay constants differed from zero within each epoch and to compare decay constants between epochs. To limit the influence of unusually large decay estimates, analyses were restricted to *k* ≤ 0.15; for comparisons between epochs, sessions were included only when *k* met this criterion in both epochs being compared.

Summary statistics are reported using nonparametric measures: median and median absolute deviation (MAD), unless otherwise noted. Differences between conditions or relative to baseline were assessed using Wilcoxon signed-rank tests. For the 8-targets task, the relationship between neural parameters (e.g., CSD value or recording depth) and target location (i.e., target proximity) was evaluated using a nonparametric circular-linear correlation test (Zar 1999). Note that, to compare sink (negative values) and source (positive values) hotspot values, all the sink values were multiplied by −1 to be positive. For each individual session, paired t-tests were performed between the sink or source values during the periods of interest (i.e., visual, motor and post-motor epochs) and the baseline values at the same depth.

## Acknowledgements

We thank members of the Gandhi and Mayo labs for scientific discussions and critical feedback; Drs. A. Fisher and V. Mrotz, the veterinary technicians and husbandry staff of the Division of Laboratory Animal Resources for assistance with animal care; and Dr. I. Smalianchuk for developing the behavioral control software used in a subset of the experiments. The study was funded by the following NIH grants: R01EY037837 and P30EY008098.

## Author Contributions

C.B. and N.J.G. conceived and designed the study. C.B. performed the experiments, analyzed the data and drafted the manuscripts. F.Y. contributed to data collection and edited the manuscript. C.B. and N.J.G wrote the manuscript.

## Code and Data Availability

The data for this manuscript are available at https://doi.org/10.5281/zenodo.24323448.

---

**Supplementary Figure 1.**
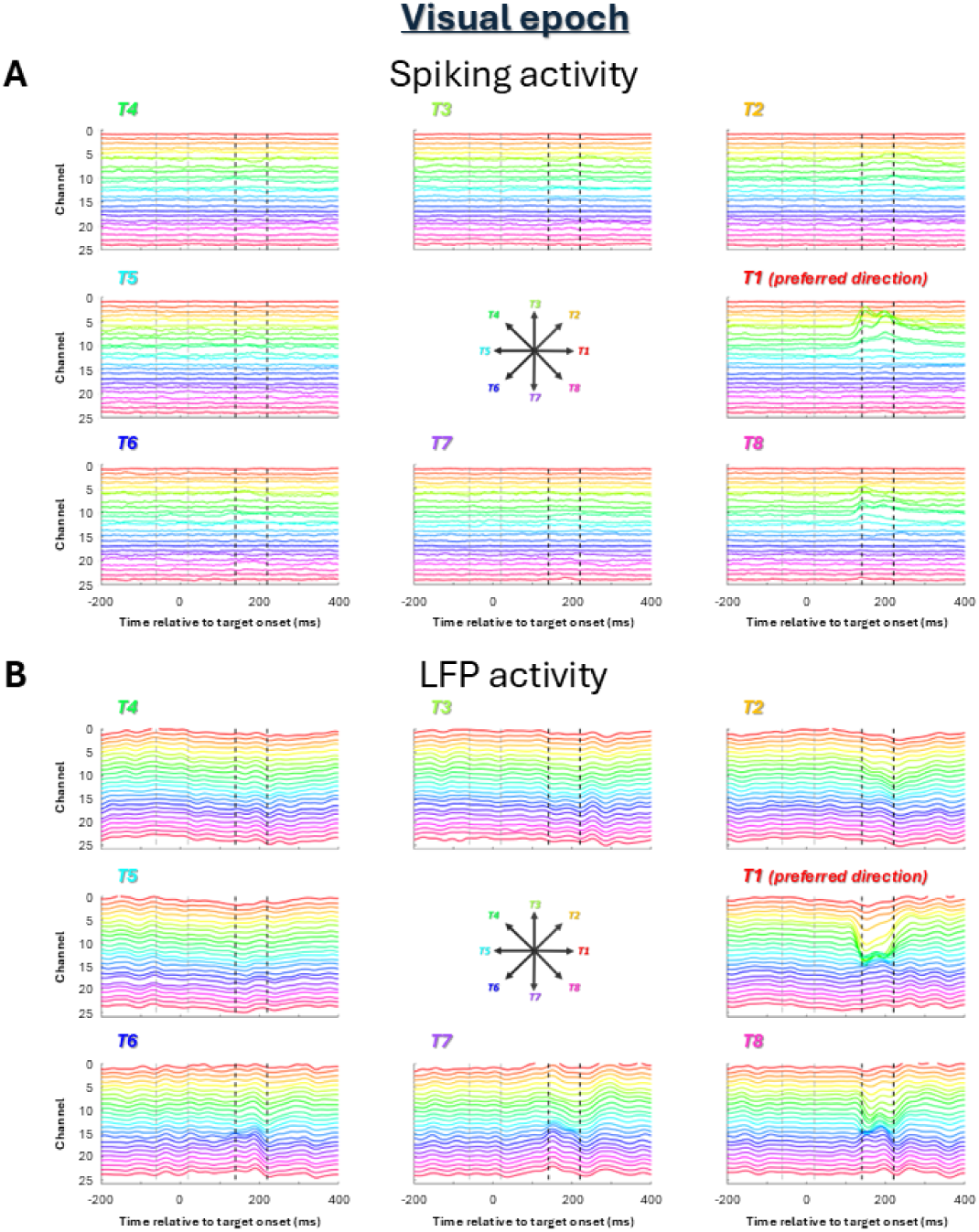
Spiking and LFP activity during the visual epoch in the eight-target task. Data are from the same example dataset shown in Figure 4. Average spiking activity (A) and LFP activity (B) across depth are shown aligned to target onset for each of the eight target locations (T1-T8). T1 corresponds to the preferred direction; the remaining targets are arranged at successive 45° angular distances around the target array. Traces are color-coded by recording channel. Dashed vertical lines indicate the visual analysis window.

**Supplementary Figure 2.**
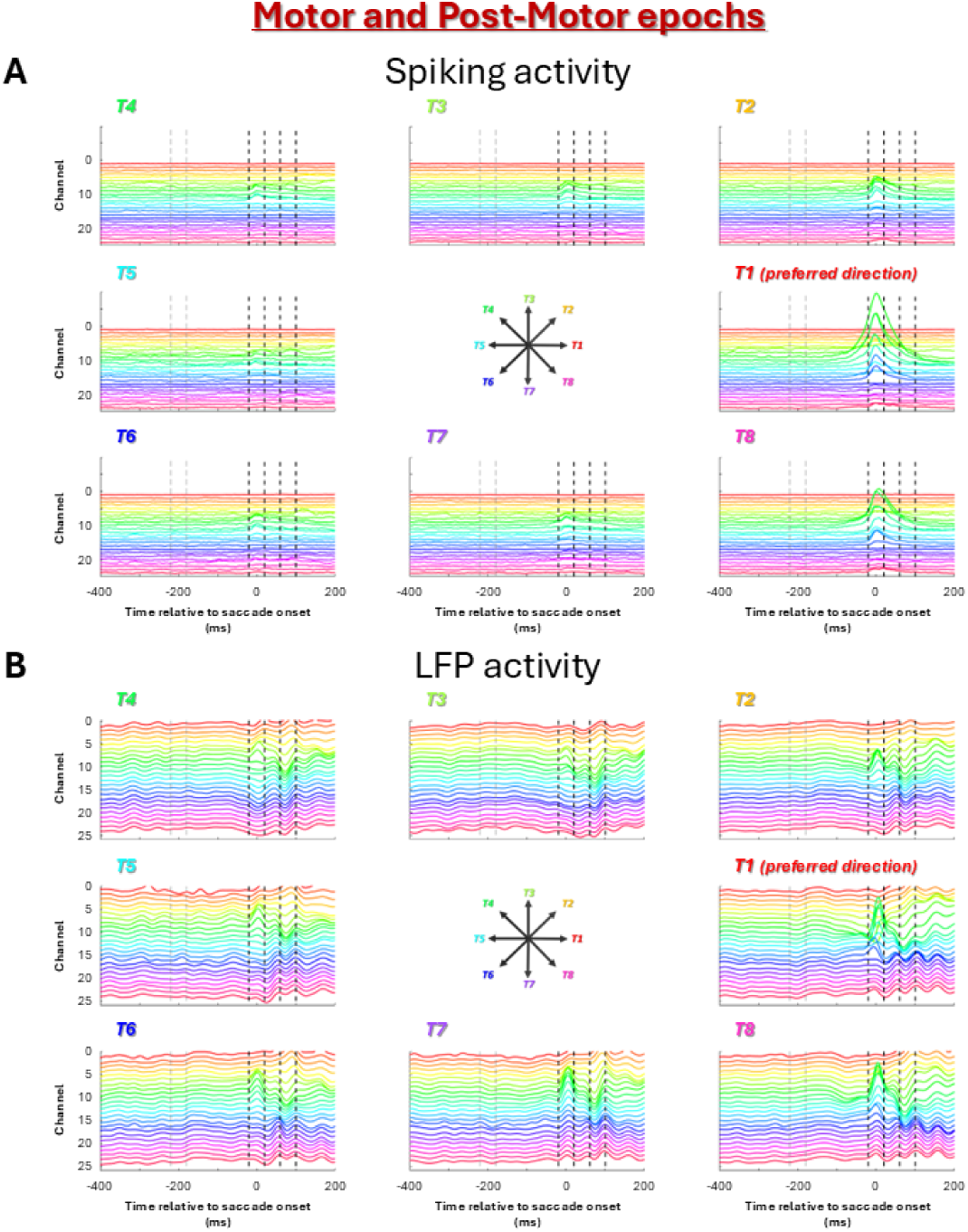
Spiking and LFP activity during the motor epoch in the eight-target task. Data are from the same example dataset shown in Figure 4. Average spiking activity (A) and LFP activity (B) across depth are shown aligned to saccade onset for each of the eight target locations (T1-T8). T1 corresponds to the preferred direction; the remaining targets are arranged at successive 45° angular distances around the target array. Traces are color-coded by recording channel. Dashed vertical lines indicate the motor analysis window.

**Supplementary Figure 3.**
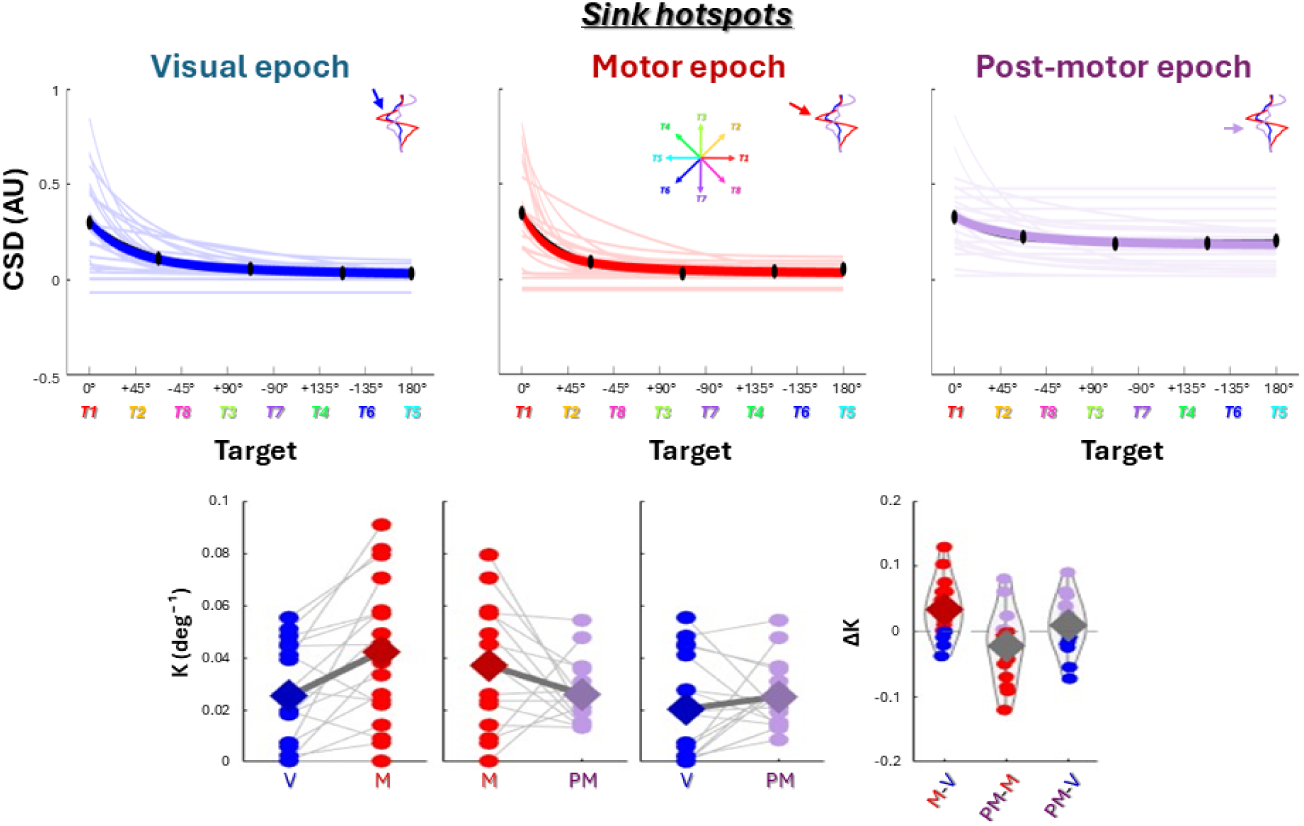
Session-wise exponential fits and decay constants for sink hotspots. Top panels show the spatial decay of sink magnitude during the visual (blue), motor (red), and post-motor (purple) epochs. For each session, sink magnitude was collapsed across targets with the same angular distance from the preferred direction (0°, 45°, 90°, 135°, and 180°) and fit with an exponential function. Thin colored lines show individual session fits, and the thick colored line represents the mean of the individual session fits. Black points show the grand-mean sink magnitude at each angular distance, and the black curve shows the exponential fit to these grand-mean values. Bottom panels compare session-wise decay constants (κ) between the visual (V), motor (M), and post-motor (PM) epochs. Analyses were restricted to κ ≤ 0.15, with filtering performed separately for each pairwise comparison such that both κ values were ≤ 0.15. Colored circles represent individual sessions, and thin gray lines connect values from the same session. Diamonds indicate the mean κ for each epoch, with thick gray lines connecting the means. The rightmost panel shows the distribution of session-wise differences in decay constants (Δκ) for M-V, PM-M, and PM-V. Colored circles represent individual session differences, diamonds indicate the mean Δκ, and the dashed horizontal line indicates zero.

**Supplementary Figure 4.**
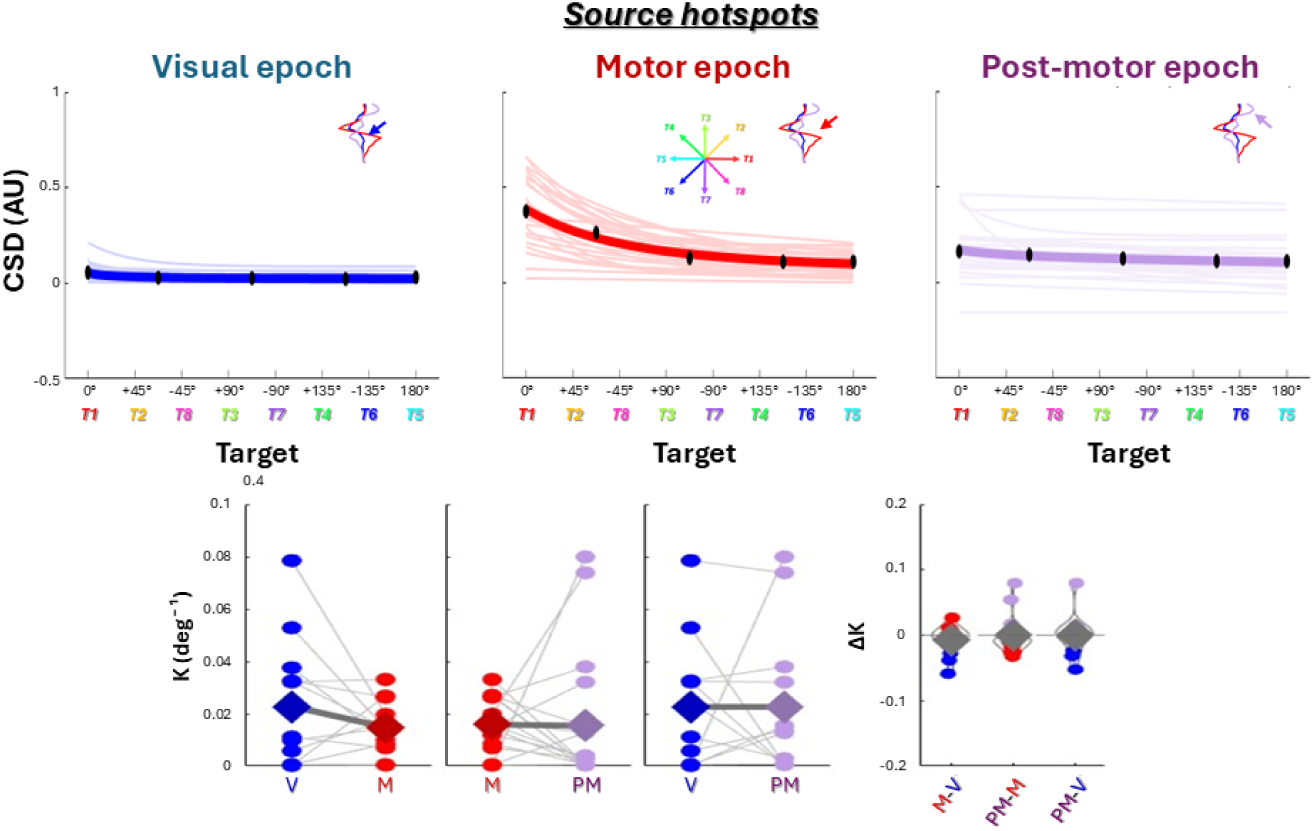
Session-wise exponential fits and decay constants for source hotspots. Top panels show the spatial decay of source magnitude during the visual (blue), motor (red), and post-motor (purple) epochs. For each session, source magnitude was collapsed across targets with the same angular distance from the preferred direction (0°, 45°, 90°, 135°, and 180°) and fit with an exponential function. Thin colored lines show individual session fits, and the thick colored line represents the mean of the individual session fits. Black points show the grand-mean source magnitude at each angular distance, and the black curve shows the exponential fit to these grand-mean values. Bottom panels compare session-wise decay constants (κ) between the visual (V), motor (M), and post-motor (PM) epochs. Analyses were restricted to κ ≤ 0.15, with filtering performed separately for each pairwise comparison such that both κ values were ≤ 0.15. Colored circles represent individual sessions, and thin gray lines connect values from the same session. Diamonds indicate the mean κ for each epoch, with thick gray lines connecting the means. The rightmost panel shows the distribution of session-wise differences in decay constants (Δκ) for M-V, PM-M, and PM-V. Colored circles represent individual session differences, diamonds indicate the mean Δκ, and the dashed horizontal line indicates zero.

